# Site-specific recognition of SARS-CoV-2 nsp1 protein with a tailored titanium dioxide nanoparticle

**DOI:** 10.1101/2021.07.27.453834

**Authors:** P. Agback, T. Agback, F. Dominguez, E.I. Frolova, G. Seisenbaeva, V. Kessler

## Abstract

The ongoing world-wide Severe Acute Respiratory Syndrome coronavirus 2 (SARS-CoV-2) pandemic shows the need for new sensing and therapeutic means against the CoV viruses. The SARS-CoV-2 nsp1 protein is important, both for replication and pathogenesis, making it an attractive target for intervention. In recent years nanoparticles have been shown to interact with peptides, ranging in size from single amino acids up to proteins. These nanoparticles can be tailor-made with specific functions and properties including bioavailability. To the best of our knowledge, in this study we show for the first time that a tailored titanium oxide nanoparticle interacts specifically with a unique site of the full-length SARS-CoV-2 nsp1 protein. This can be developed potentially into a tool for selective control of viral protein functions.

## Introduction

SARS-CoV-2 is a member of the *Betacoronavirus* (β-CoV) genus. Its genome is represented by a single-stranded RNA of ∼30 kb in length^1, 2, 3^. The genomic RNA is directly translated into two very long polyproteins, which are further processed into the individual non-structural proteins nsp1 to nsp16 by the encoded protease activities. These proteins are viral components of the replication complexes, and also induce modification in the intracellular environment required for efficient viral replication. Nsp1 protein of coronaviruses is the major determinant of viral pathogenesis^4, 5, 6^. It is known to efficiently inhibit cellular translation, induces degradation of cellular mRNAs and, thus, prevents the development of antiviral response^7, 8, 9, 10, 11, 12, 13, 14, 15, 16, 17, 18, 19^. Nsp1 is also required for viral replication, buts its role remains elusive. This makes it an attractive target for development of therapeutic means against SARS-CoV-2 infection. Nsp1 is a relatively small (20 kDa) and contains a folded domain (aa 10 to 124) and two intrinsically disordered domains. The last 26-aa-long fragment of nsp1 is responsible for binding with the 40S ribosome subunit. Function and interactive partner of the folded domain remains unknown. Recently, structural analysis of SARS-CoV-2 nsp1 has been performed by our and other groups. Particularly, two X-ray structures of the folded domain (7K7P and 7K3N), and secondary structure determination of the full-length nsp1 based on NMR assignment of the backbone resonances have been published^20, 21, 22, 23^. In our study, we have also made partial side-chain assignment^23^. Availability of the structural information allows now to analyze nsp1 interaction with cellular and viral proteins and small molecules.

Metal oxide nanoparticles (NP), *i*.*e*., nanoparticles of common sand minerals, are abundant in ground water and watercourses, due to both physical and chemical erosion via dissolution and re-crystallization of ground minerals^24^. Their surface can be rather chemically reactive, and are known to be able to produce or destroy reactive oxygen species by catalyzing RedOx reactions, especially in daylight via photochemical mechanisms^25^. On the other hand, they are also known to interact strongly and specifically with biomolecules, especially proteins. Pristine NP are practically unknown in biological systems, because of the “protein corona” phenomenon, aiming to describe the assembly of protein molecules on the surface of larger NP. Smaller NP are suspected to interact specifically with selected fragments in the structure of *e*.*g*., blood proteins^26^. The nature of proteins, specifically adsorbed by NP is related not only to their size, but, most importantly, to the chemical composition of the NP. The effects of interactions may be drastically different. Some NP, like those of alumina, Al_2_O_3_, Boehmite, AlO(OH), or zirconia, ZrO_2_, have been reported to catalyze hydrolysis of proteins, while other, like TiO_2_, were proven to cause protein coagulation^27, 28, 29^. Molecular interactions between NP and proteins have until recently been very scarcely characterized. It has been noticed that proteins and NP can co-assemble into micro-sized colloid crystals^30^. An attractive possibility to investigate such interactions would be the studies of complexes derived from individual poly oxo-metalate species (POMs). Importance of possible specific POM-protein interactions for potential biological applications has been outlined in a recent review^31^. POMs can be considered as individual molecular species but are at the same time nanoparticles with a size just above 1 nm. Until recently, only very few structures of proteins with POMs were reported, suffering of rather poor resolution of the area surrounding often strongly disordered POM species^32^.

In this study we have applied nuclear magnetic resonance (NMR) spectroscopy in combination with tuning of the chemical composition and surface structure of titanium oxide nanoparticles (NP) to probe their affinity to full-length of SARS-CoV-2 nsp1 protein. Our on-going study of the structure of protein and nanoparticle complexes will reveal the mechanism underlying this process and can be developed into a tool for selective control of viral enzymes.

## Results

For investigation of interaction with SARS-CoV-2 nsp1 protein we selected two kinds of well-characterized small Titanium dioxide nanoparticles, possessing anatase structure. One of them NP1 with hydrodynamic diameter 3.8 nm and zeta-potential − 8.2 mV was obtained from industrially produced TiBALDH precursor and was proved to be terminated by lactate ligands^33^. The other material NP2 was produced from triethanolamine modified precursor by acidic hydrolysis and proved to contain particles with hydrodynamic diameter 3.2 nm and zeta-potential − 11.4 mV^34^. The measured negative potential was indicating that the ligand was in aqueous medium essentially desorbed from the particles, leaving them with pristine, supposedly, partially hydroxyl terminated surface. The zeta-potential of this second type of titania, − 11.4 mV, was quite close to that of sol-gel TiO_2_ obtained by just hydrolysis with water of pure alkoxide precursor with subsequent peptization and equilibration with water, − 11.7 mV^35^. The behavior of the particles upon titration with the protein turned rather different: while addition of NP1 in 1:1 molar ratio did not produce any measurable effect on the 1H-15N TROSY NMR spectrum of nsp1, that of NP2 resulted in very distinct signal perturbation for several amide resonances of the second half of the rigid α-helix fragment of nsp1 (L39, E41 – G49) as well as that of H110 (Fig 1). A further addition of NP1 to the sample to increase the ratio to 10:1 did not result in any changes as well. Addition of NP2, to increase the ratio to 2:1 resulted in only minor changes for the amides mentioned earlier. The results clearly show site specific binding of NP2 to the α-helix of nsp1.

**Figure 1.**
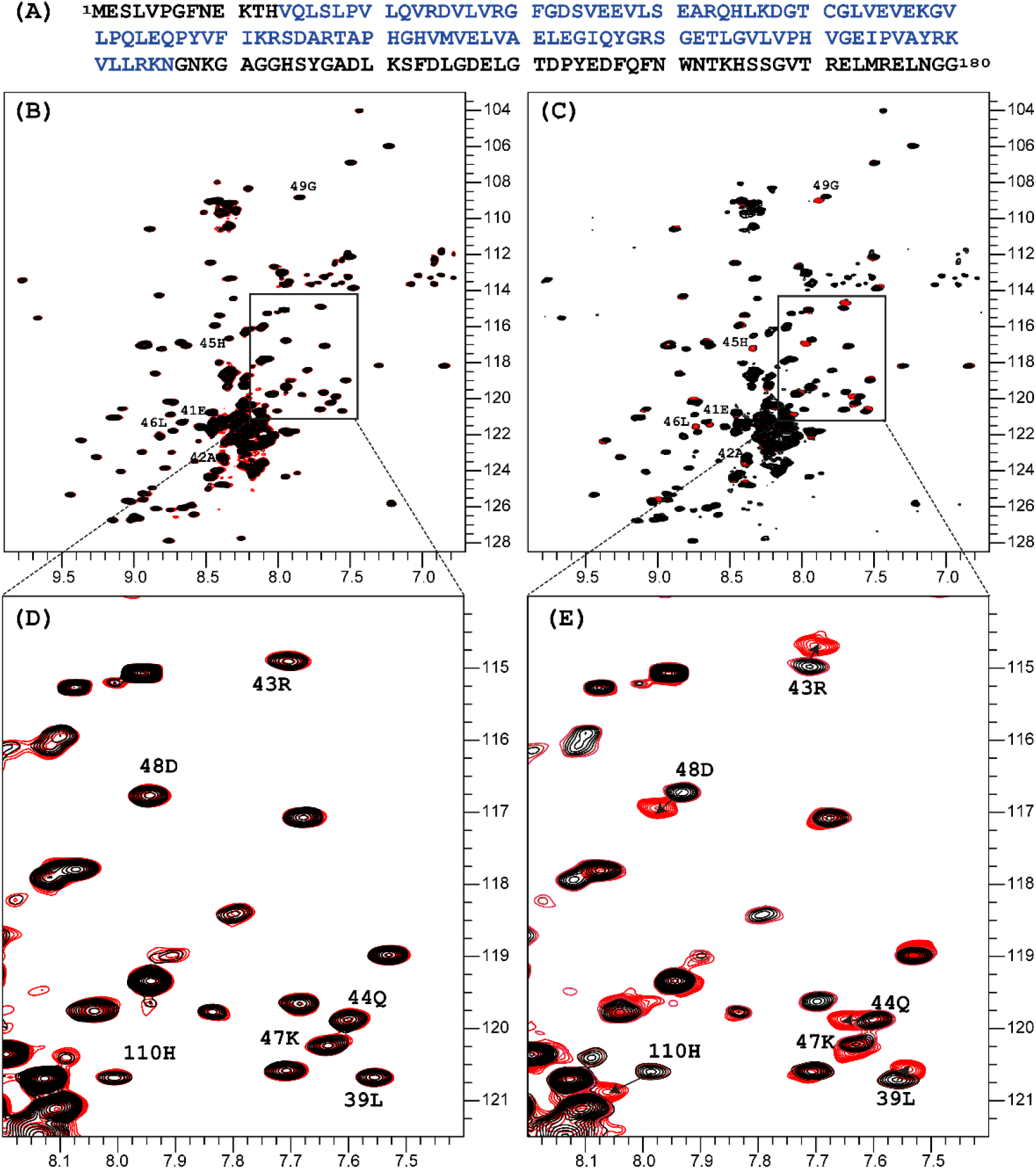
Sequence of the full length of SARS-CoV-2 nsp1 protein is presented on top in panel, in blue colour is the sequence used in crystallisation 7K7P (**A**). Superposition of the ^1^H-^15^N TROSY spectra of SARS-CoV-2 nsp1 protein apo form (black colour) and apo with added nano particles (red colour) are shown in panels (**B**)-(**E**). For the added NP1 particle, ratio nsp1 to NP1 is 1 to 10, spectra presented in (**B**) and its expanded part in (**D**) panels are identical to the apo form of nsp1 indicating that there is no interaction between NP1 and nsp1. For the added NP2 particle, ratio nsp1 to NP2 is 1 to 2, spectra presented in (**C**) and its expanded part in (**E**) panels are different. The chemical shift assignment of NH backbone resonances with induced chemical shift perturbation by binding of NP2 to nsp1 are shown by the number and symbols corresponding to the sequence (**A**).

This difference between the NPs originated most probably from the fact that the surface of NP1 was blocked by strongly bound lactate ligands, while that of NP2 remained highly reactive. The major driving force in the interaction of NP2 with the nsp1 protein was most likely the formation of inner sphere carboxylate complexes on the surface achieved by binding of the side chain carboxylate groups of aspartic and glutamic acids. The structure of such complexes always involves binding of the two oxygen atoms of the carboxylate groups to two adjacent Ti atoms connected via a double oxygen bridge Ti(μ -O)_2_Ti (see Fig. 2a). The attachment of the carboxylic group appears quite strong and is surprisingly uniform with T-O distances ca. 2.05 and ca. 2.10 Å respectively^36^.

**Figure 2.**
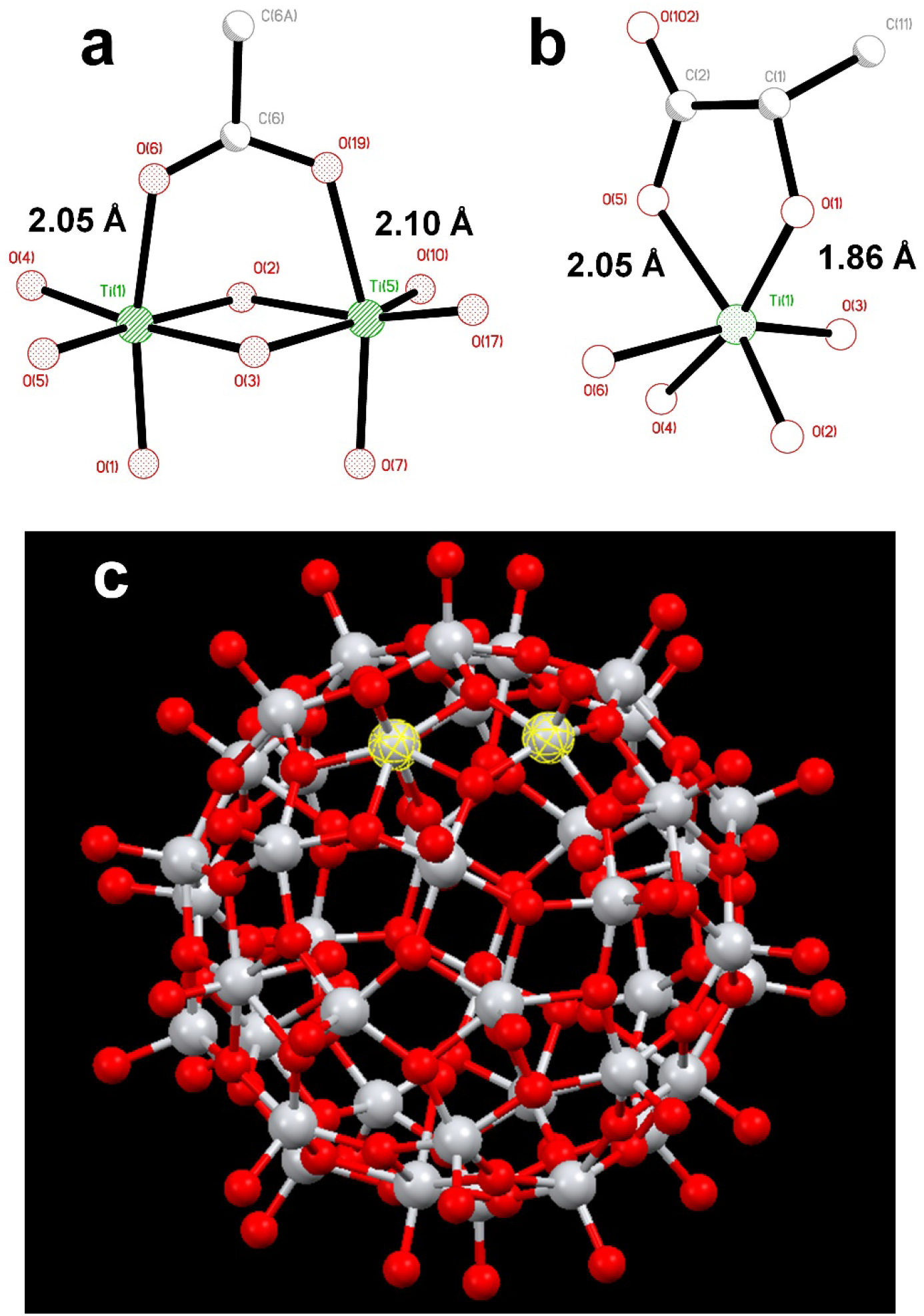
Molecular structures of (a) fragment, demonstrating the binding of a carboxylate ligand to TiO_2_ (anatase type) surface^36^; (b) fragment showing commonly observed chelated bonding of a lactate ligand to a Ti atom^33^; (c) the reported titanium-oxygen core in the structure of H_6_[Ti_42_(μ_3_-O)_60_(O^i^Pr)_42_(OH)_12_)] structure^37^. A Ti(μ -O)_2_Ti fragment on the surface capable to binding a carboxylate group is highlighted.

This binding appears, however, to be substantially weaker than that of a lactate ligand. Lactate units, as revealed by the literature data, and the study of the new compound reported in this work, are always bound in a chelating mode to a single titanium atom^33^. It features a Ti-O distance to a single-bound oxygen atom in the carboxylate typical for such fragments, ca. 2.05 Å (see Fig. 2a), and with a much shorter (and thus stronger) Ti-O bond to the alkoxide oxygen in the lactate structure of ca. 1.86 Å (see Fig. 2b). Inner sphere bonding corresponding to the mode typical for carboxylate binding is thus possible to a pair of adjacent Ti atoms connected via a pair of O-bridges as indicated in Fig. 2c. This may provide explanation for the absence of interaction between lactate-terminated TiO_2_ and the protein. Still the binding of NP2 is complete, indicated by the fact that further addition did not induce any further chemical shift perturbations. Most likely other interactions than Ti-O are involved.

In order to provide closer molecular insight into the interaction of the NP2 material with SARS-CoV-2 nsp1 protein, we decided to investigate possibility of geometrical matching between a model TiO2 particle and the structure of the SARS-CoV-2 nsp1 protein reported from the X-ray single crystal study (7K7P)^21^. To represent the curved anatase structure we used the model of spherically shaped anatase monolayer in the recently reported H_6_[Ti_42_(μ_3_-O)_60_(O^i^Pr)_42_(OH)_12_)] structure^37^. The cif-file was obtained from the supplementary materials to the publication and was lacking both carbon and hydrogen atom positions making it ideal for geometrical matching by using the program Chimera^38^. It was converted into PDB-format and used for docking. What we observe in the 1H-15N TROSY spectra is of course chemical shift changes of the backbone amide groups. It is very unlikely that of the amides would directly interact with the nanoparticle without major structural changes of nsp1. Most likely it is the sidechains that interacts with the titanium-oxide nanoparticle. One of possible docking configurations consistent with the NMR spectrum in Fig. 1 is displayed in Fig. 3. The conclusion that could be obtained from matching the surface geometries was that the TiO_2_ particle was experiencing a specific attraction resulting in inner sphere complex formation with one carboxylate fragment, originating from D48 and additional possible hydrogen bonding to the sidechains of E41 (carboxylate), Q44 (amide carbonyl) and H45 (Nδ1). The role of H110 is unclear, further studies are necessary to understand its role. The results of geometric matching are in good agreement with the observed changes in the NMR spectrum. This indicates that the structure of H_6_[Ti_42_(μ_3_-O)_60_(O^i^Pr)_42_(OH)_12_)] has a good resemblance with the structure of small anatase NP. One should note that the NP2 used in this study is larger than the nanoparticle used in the modelling with a presumably flatter surface making more interactions with the α-helix possible.

**Figure 3.**
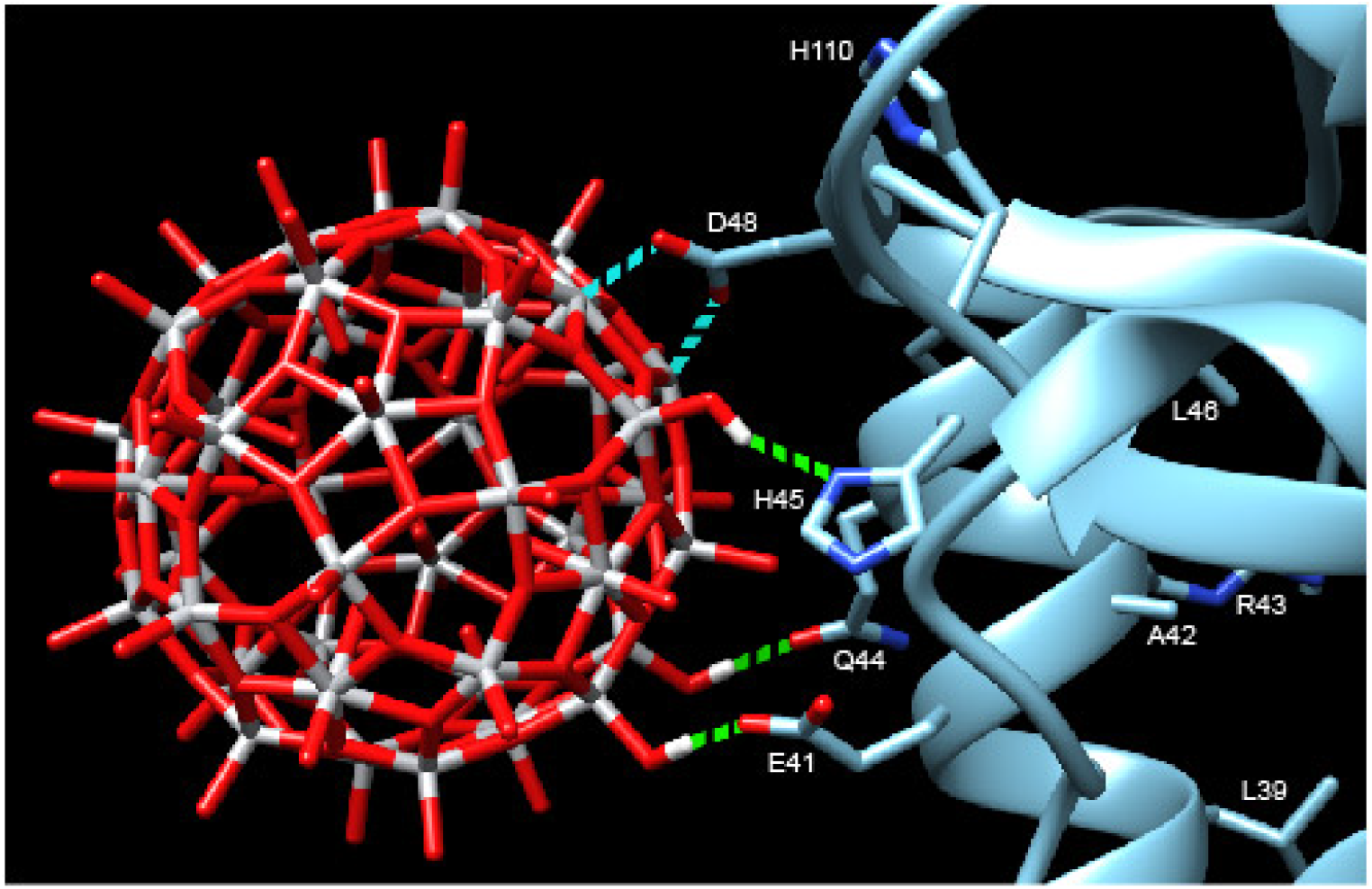
Model showing the docking of the metal-oxygen core of H_6_[Ti_42_(μ_3_-O)_60_(O^i^Pr)_42_(OH)_12_)] with the structure of nsp1 taken from 7K7P^21, 37^. The possible interactions are show in cyan: carboxylate of D48 with titanium, forming an inner sphere complex, and green: hydrogen bonding of OH-groups from the nanoparticle to E41, Q44 and H45. The amino acids of the other amides showing chemical shift perturbation are marked.

## Discussion

The full-length nsp1 protein contains several structural elements: one α-helix, β-sheets, turns and one short and one long disordered domain. It is not fully clear as to why we only see site specific binding to the second half of the α-helix. There are 12 glutamic acids and 4 aspartic acids in the sequence, all presumably possible interaction points, if one assumes that the carboxylate binding is the main driving force. Most likely the distance between them and the possibility of other hydrogen bond partners are a good fit for the surface of the titanium oxide nanoparticle. It is not clear as to why H110 shows significant perturbation as well. The sidechain seems to be too far away for hydrogen bonding, but it might possibly be affected by changes in stacking interaction with H45 induced by the nanoparticle binding. Further investigations of the nature of this interaction are being planned. The studies will necessarily have to interweave the disciplines of inorganic chemistry and structural biology which has been considered to hold great interest in the future. As a starting point we are now looking to map the changes in the sidechains of the nsp1 protein upon nanoparticle binding by NMR spectroscopy. By this we hope to precisely define the nature of the interaction between the titanium oxide nanoparticle and the nsp1 protein. More detailed docking studies are also necessary to understand the interactions between the nanoparticle and the nsp1 protein. The calculation by either quantum or molecular mechanics of a protein structure together with a metal-oxide nanoparticle will be a challenging task.

The possibility of specific geometrical matching with resulting strong binding between NP and the protein, opens the possibility for chemical tuning of NPs. This includes modifying the chemical composition and the surface structure of the NPs. The goal is then to make them a potential tool for blocking protein recognition sites with a broad area of possible applications ranging from new techniques of rapid detection of viral antigens to creating new anti-viral drugs.

It should be mentioned that antiviral activity of metal oxides has been traced recently in a number of cases and an effect of complex cerium oxide NP specifically against SARS-Cov-2 was observed^39, 40^. However, no mechanism of action for this have been suggested yet. Our reported finding in this study could provide a possible mechanism for this effect.

This finding also shows that nanoparticles could be used as a tool for the mapping of the structure-dynamic landscape of active proteins, especially if they could be tailormade to find specific structural motifs.

## Acknowledgments

The authors would like to express their gratitude to Swedish Research Council (Vetenskapsrådet) for support of the Grant No. 2018-03811, Molecular mechanisms in interaction of mineral nanoparticles and proteins. This work was also supported by the Swedish Foundation for Strategic Research, Grant ITM17-0218, Innovative Experimental Modeling of Dynamic Protein States. EIF acknowledges the support of NIH grant R21AI146969 and UAB Research Acceleration Funds.

## Materials and methods

### Titania nanoparticles

Concentrated solution of TiBALDH, 50 wt% with respect to (NH_4_)_8_Ti_4_O_4_(OCOCHOCH_3_)_8_·4H_2_O was obtained from Sigma Aldrich and used for producing lactate-capped TiO_2_ NP by dilution with MilliQ water. The starting concentration of TiO_2_ was estimated according to established solution equilibrium as 45 mg/ml^33^. The starting solution of triethanolammonium capped TiO_2_ NP was obtained following the procedure described in reference 36 by dissolving Ti(OEt)_4_ (5 mL) in anhydrous EtOH (5 mL), adding first 1.5 mL of triethanolamine and then 1 ml of hydrolyzing solution, prepared by mixing 0.5 M HNO_3_ (0.5 mL) with EtOH (2.0 mL). This procedure was resulting in a starting solution with concentration of TiO_2_ NP equal to 120 mg/g as confirmed earlier by TGA. Dilution was carried out with MilliQ water. Formal molar concentrations of TiO_2_ NP were calculated by dividing the TiO_2_ mass content in solution by estimated mass of a single particle, assuming the hydrodynamic sizes of lactate-capped TiO_2_ NP as spheres 3.8 nm and triethanolammonium capped TiO_2_ NP as 3.2 nm in diameter respectively^33, 41^. Both kinds of NP have earlier been proved to possess anatase structures.

As model for surface binding was chosen the structurally characterized and chemically individual spherical particle reported in 1.55 nm in geometric diameter^37^.

[(NH_4_)_2_Ti(OCOCHOCH_3_)_3_]_2_·EtOH(**1**). Mother liquor after sedimentation of lactate-modified TiO_2_ particles by addition of equal amount of 96% ethanol to 50 wt% solution of TiBALDH in water was left for drying in air, producing the rod-shaped crystals of **1**.

### Characterization

TiO2 content of the triethanolamine-derived sols was determined with PerkinElmer Pyris 1 thermobalance.

### Crystallography

The data collection has been carried out at room temperature (23±2°C) using Bruker D8 SMART Apex2 CCD diffractometer with MoK_α_ radiation (λ= 0.7083 Å) in the 2θ range 6.28 – 50.04°. C_20_H_50_N_4_O_15_Ti_2_, M= 682.44 Da, Monoclinic, Space Group C2, *a* = 15.053(9), *b* = 8.727(5), *c* = 25.957(15)Å, β = 91.475(10)°, V = 3409(3) Å^3^, d_calc_ = 1.330 g/cm^3^. 5760 symmetrically independent reflections with R_int_ = 0.0346 were obtained from data integration. The structure was solved by direct methods. The coordinates of all non-hydrogen were obtained from the initial solution and refined in isotropic and then anisotropic approximation. The hydrogen atoms were added by geometrical calculation for C-atoms and by differential Fourier syntheses for N-atoms and refined isotropically using a riding model. Final discrepancy values were R1 = 0.0362, wR2 = 0.0833 for 5098 reflections with I > 2sigma(I)). Full details of structure solution and refinement are available free-of-charge from the Cambridge Crystallographic Data Centre citing the reference number **2086690**, at https://www.ccdc.cam.ac.uk/solutions/csd-core/components/csd/.

### Protein expression and purification

The protein was prepared as described in reference 25.

### NMR samples preparation

All NMR experiments were performed in a buffer containing 20 mM HEPES pH 7.5, 1 mM NaN_3_, 10 (v/v) % D_2_O and 0.1 mM DSS (4,4-dimethyl-4-silapentane-1-sulfonic acid) as an internal ^1^H chemical shift standard. Two such samples were prepared. The protein concentration was 0.25 mM, and all spectra were acquired in 3 mm tubes (final volume of 0.2mL). ^13^C and ^15^N chemical shifts were referenced indirectly to the ^1^H standard using a conversion factor derived from the ratio of NMR frequencies. The nanoparticles, NP1 and NP2, were dissolved in H2O for a concentration of 500 mM and 50 mM, respectively.

### NMR experiments

NMR experiments were acquired on Bruker Avance III spectrometers operating at 14.1 T, equipped with a cryo-enhanced QCI-P probe. All ^1^H-^15^N TROSY experiments were run with 400 increments and 16 scans. As the first step we recorded TROSY spectra for the protein without anything added. Then we added 1 uL of the respective NP solution to form 1:1 complexes (NP1 solution was diluted 10 times with D2O before) and TROSY was ran again. In a final step, 1 ul of NP1 was added to its 1:1 complex to change it into 10:1 and 1ul of NP2 to its 1:1 complex to increase it to 2:1. Again TROSY spectra were recorded.

### Modelling

Molecular graphics and analyses was performed with UCSF Chimera, developed by the Resource for Biocomputing, Visualization, and Informatics at the University of California, San Francisco, with support from NIH P41-GM103311. The 7K7P pdb file as well as a pdb file for the nanoparticle obtained from a cif file of the H_6_[Ti_42_(μ_3_-O)_60_(O^i^Pr)_42_(OH)_12_)] structure was combined in chimera 1.15 and docking could be obtained by matching the sidechains of the amino acids showing chemical shift perturbation with the surface of the nanoparticle^38^.

